# Beyond harm: synthesising meta-analytical and mechanistic evidence on animal responses to microplastics

**DOI:** 10.64898/2026.09.19.752891

**Authors:** Colette Martin, Pablo Burraco, Katharina Ruthsatz

## Abstract

A central question in ecology and evolution is determining when environmental challenges translate into stress and biological harm. Human-derived pollutants are often assumed to be inherently damaging; however, their effects may depend strongly on ecological and physiological context. Microplastics exemplify this complexity, eliciting heterogeneous responses across taxa rather than uniform toxicity. Here, we synthesise evidence from 14 meta-analyses encompassing 9,101 effect sizes to identify the conditions under which microplastic exposure leads to damage, neutrality, compensation, or stimulation. These syntheses reveal clear negative effects alongside frequent neutral outcomes and occasional stimulatory responses. Microplastic effects vary with particle characteristics and exposure conditions and are strongly moderated by species-specific traits and life stage. We further argue that plasticity in the gut and in behavioural and physiological traits may buffer microplastic effects. Emerging evidence also indicates that biofilm colonisation alters the physicochemical and biological properties of microplastics, potentially modifying their ecological interactions and toxicity profiles. By reframing microplastics as context-dependent challenges rather than universally toxic agents, we provide a conceptual framework for understanding animal responses under realistic ecological conditions and identifying the biological and environmental contexts in which microplastic exposure is most likely to result in harm.

## 1. Introduction

Global change is exposing organisms to novel and complex environments, including a growing diversity of human-derived pollutants (Bernhardt et al., 2017; Pereira et al., 2012; Singh et al., 2023). Pollution is often treated as a pervasive source of physiological disruption, with the potential to divert energy from growth and reproduction towards somatic maintenance (Brander et al., 2026; Jager & Zimmer, 2012). However, exposure alone does not necessarily translate into biological harm: outcomes can range from reduced performance to neutral, compensatory, or even stimulatory responses, depending on ecological context, exposure conditions, and the capacity of organisms to buffer environmental stress (Hossain & Olden, 2022; Jager & Zimmer, 2012; Wang et al., 2024). Understanding this variation is essential for predicting when pollutants become ecologically and evolutionarily relevant stressors rather than merely environmental exposure (Burraco et al., 2025).

Microplastics, defined as plastic particles smaller than 5 mm (Frias & Nash, 2019), have become emblematic pollutants of the Anthropocene, now widespread across terrestrial and aquatic ecosystems, trophic interactions, and food webs worldwide (Auta et al., 2017; Rillig & Lehmann, 2020; Thompson et al., 2004). Yet microplastics are not a single, uniform stressor. They originate both from primary sources, where particles are manufactured at microscopic sizes, and the fragmentation of larger plastic debris, generating a highly diverse pool of particles that differ in size, shape, polymer type, chemical composition, ageing state, and biological colonisation (Andrady, 2015; Murphy et al., 2016). This heterogeneity makes microplastics a particularly powerful system for asking when anthropogenic exposure becomes physiological stress, and when organisms instead buffer, compensate for, or tolerate such exposure without measurable harm.

Microplastics often enter animals via ingestion, respiratory pathways, or direct contact with external tissues such as eyes and skin (Mansfield et al., 2024; Rahman et al., 2024; Wang et al., 2024). Ingested particles can accumulate within the digestive tract and translocate across the intestinal epithelium, reaching secondary organs and potentially triggering systemic effects (Dong et al., 2023). For instance, the accumulation of microplastics within the digestive tract may induce a false sense of satiation or reduce energy intake, thereby affecting growth and behaviour (Cole et al., 2013; Hossain & Olden, 2022; Wang et al., 2024). Microplastic exposure can also induce physiological stress, including oxidative stress and immune suppression (Abbas et al., 2025; Bao et al., 2025; Rochman et al., 2013; Wang et al., 2024). Such effects have understandably reinforced a damage-centred view of microplastic pollution.

The absence of measurable harm, however, does not necessarily imply biological irrelevance. Responses to microplastic exposure seem to be heterogeneous across taxa, life stages, exposure regimes, and response traits (Hossain & Olden, 2022; Moyo, 2022; Wang et al., 2024). Some organisms show little measurable change, whereas others maintain performance despite physiological activation, or display responses consistent with compensatory plasticity or hormesis. These outcomes are important because they reveal when organisms can buffer pollutant exposure, which mechanisms may underlie apparent resilience and where the limits of compensation are likely to emerge. Despite this variability, most research has primarily emphasised detrimental outcomes, leaving neutral, compensatory, and potentially stimulatory responses comparatively underdeveloped in ecological and evolutionary interpretations of microplastic effects.

In this integrative synthesis, we move beyond a damage-centred perspective to examine when and why microplastic exposure does not translate into harm. First, we examine evidence from meta-analyses to identify the ecological and physiological conditions under which microplastic exposure results in negative, neutral, compensatory, or stimulatory outcomes across animals. Second, we evaluate how exposure properties, including particle characteristics, dose, and duration, interact with species-specific traits and physiological plasticity to shape these outcomes. Third, we discuss how compensatory responses in behaviour, energy allocation, antioxidant regulation, immunity, and organ plasticity may buffer microplastic exposure, but may also generate hidden or delayed effects. By integrating these processes within an ecological and evolutionary framework, we highlight their implications for organismal resilience, population persistence, and conservation in environments increasingly polluted by microplastics.

## 2. Meta-analytical evidence for harm, neutrality, and stimulation

Quantitative syntheses and meta-analyses evaluating the effects of microplastics on animal health and physiology now provide a broad evidence base rather than isolated case studies. To summarise this literature, we searched Scopus, Web of Science, and PubMed on 29^th^ May 2026 using the search string [( “microplastic*” AND ( “health” OR “fitness” OR “performance” OR “effect*” ) AND ( “mammal*” OR “fish*” OR “amphibian*” OR “cladocerans” OR “reptile*” OR “bird*” OR “vertebrat*” OR “Protozoa” OR “Placozoa” OR “Porifera” OR “Cnidaria” OR “Ctenophora” OR “Platyhelminthes” OR “Nematoda” OR “Rotifera” OR “Annelida” OR “Echinodermata” OR “Hemichordata” OR “Mollusca” OR “Gastropoda” OR “Bivalvia” OR “Cephalopoda” OR “Arthropoda” OR “Insecta” OR “Crustacea” OR “invertebrat*” OR “marine organism*” OR “freshwater organism*” OR “terrestrial organism*” ) AND ( “meta-analys*” OR “meta analys* “))].

The database search identified 202 records, of which 101 records remained for title and abstract screening after duplicate removal (Figure S1). Of these, 69 records were retained for full-text assessment, and 14 quantitative syntheses met the criteria for inclusion (Figure S1). Reports were eligible if they 1) evaluated microplastic exposure as a single stressor, 2) reported a quantitative measure of effect size (e.g., Hedges’ g, lnRR, odds ratio (OR), or mean difference (MD)), and 3) reported the direction and magnitude of microplastic effects on biological traits related to animal health, physiology, performance, or fitness. Where response ratios (RR) were reported, they were converted to natural log response ratios (lnRR) using ln(RR) to ensure consistency across studies reporting this effect size. For each synthesis, we extracted information on taxonomic group, response trait, exposure context, particle characteristics, and effect direction and magnitude.

Collectively, the included syntheses reported a total of 9,101 effect-size estimates (k) across vertebrates (k = 5,511) and invertebrates from aquatic (k = 2,253) and terrestrial systems (k = 1,337). These estimates encompass responses ranging from molecular and physiological endpoints to whole-organismal traits. To facilitate comparisons among meta-analyses, the response traits extracted from each synthesis were classified into 10 broad categories: behaviour and locomotion (e.g., activity, movement, swimming performance, exploration, and foraging), energy and metabolism (e.g., energy consumption, oxygen consumption, digestion, nutrient assimilation, and metabolism-related biomarkers), feeding and ingestion (e.g., microplastic ingestion and egestion, digestive processing, microplastic accumulation), growth and development (e.g., body size, growth, developmental time, and developmental abnormalities), immunity (e.g., immune function and inflammatory responses), microbiome (e.g., gut microbial diversity and composition), morphology (e.g., external and interior morphology, organ morphology, deformities, and histopathology), physiology and health (e.g., oxidative stress, neurophysiology, and detoxification pathways), reproduction (e.g., fecundity, hatching success, and reproductive hormones), and survival (e.g., survival, mortality, and lifespan) (Table S1). Detailed descriptions of the response traits extracted from each meta-analysis are provided in Supplementary Table 1.

### 2.1 Vertebrates

In vertebrates, meta-analytical evidence on microplastic effects remains largely restricted to fish, covering 62 species and 5,511 effect sizes across multiple traits linked to health and performance (Bao et al., 2025; Berlino et al., 2023; Hossain & Olden, 2022; Moyo, 2022; Sacco et al., 2026; Salerno et al., 2021; Wang et al., 2024; F. Yuan et al., 2023; X. Yuan et al., 2025). Across these syntheses, negative mean effect sizes are evident for several functional trait categories such as behaviour and locomotion (Figure 1A: Hedges’ g = -0.59 to -0.21; Figure S2A: lnRR = -0.33; Figure S3A: OR = 8.34), as well as changes in energy and metabolism (Figure 1B: Hedges’ g = -0.61 to -0.17), feeding and particle processing (Figure 1C: Hedges’ g = -1.39 to -0.14; Figure S2B: lnRR = -0.91; Figure S3B: OR = 1.65), gut microbiome diversity (Figure 1F: Hedges’ g = -0.2), morphology (Figure 1G: Hedges g’ = - 1.022 to -0.16; Figure S2E: lnRR = 0.18), and physiology and health (Figure 1H: Hedges g’ = -0.7 to 3.52; Figure S2F: lnRR = -0.11). Overall, these results indicate that microplastic exposure can disrupt key life-history and physiological axes. However, although many syntheses report negative mean effect sizes, these estimates are often small, statistically non-significant, or accompanied by substantial heterogeneity. This pattern suggests that microplastic effects in fish are conditional rather than uniform, with their magnitude and direction depending on the endpoint measured, the species considered, and the exposure scenario. In addition, the mean effect sizes of multiple functional trait categories range from negative, neutral, or positive, including growth and development (Figure 1D: Hedges’ g = - 1.54 to 0.083; Figure S2C: lnRR = -0.09; Figure S3C: OR = 9.26), immunity (Figure 1E: Hedges’ g = 0.72 to 0.16; Figure S2D: lnRR = 0.01), reproduction (Figure 1I: Hedges g’ = - 1.27 to -0.1; Figure S2G: lnRR = -0.09 to -0.08; Figure S3D: OR = 1.27 to 2.46), and survival (Figure 1J: Hedges g’ = -0.75 to 0.39; Figure S2H: lnRR = -0.04; Figure S3E: OR = 2.7).

**Figure 1.**
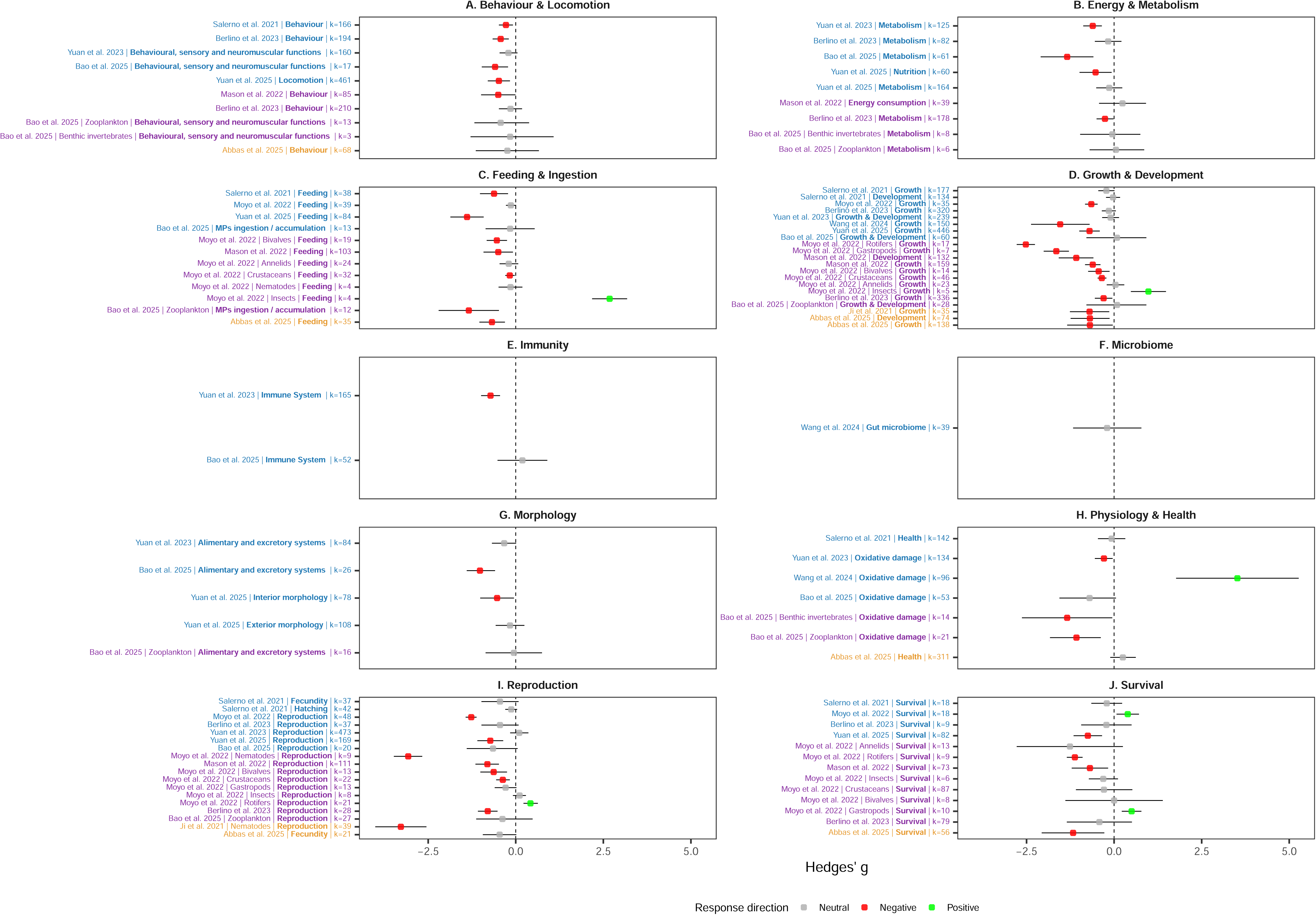
A summary of Hedges’ g effect sizes across 10 of the 14 meta-analyses, comprising vertebrates (k = 5,155), aquatic invertebrates (k = 2,095), and terrestrial invertebrates (k = 777). Hedges’ g effect sizes ±95% CI for 10 biological response trait categories under plastic exposure: A) behaviour & locomotion, B) energy & metabolism, C) feeding & ingestion, D) growth & development, E) immunity, F) microbiome, G) morphology, H) physiology & health, I) reproduction, and J) survival (detailed descriptions of the biological response trait categories are described in Table S1). Circles represent Hedges’ g effect sizes, and error bars represent 95% confidence intervals (CI). The dashed vertical line indicates no effect (g = 0), and confidence intervals overlapping zero indicate non-significant effects. The y-axis shows the meta-analysis reference, the specific response variable extracted from each meta-analysis, and the number of effect-size estimates (k). Taxonomic groups are indicated by the colour of the y-axis labels: fish (blue), aquatic invertebrates (purple), and terrestrial invertebrates (orange). Point colours indicate the biological interpretation of the response relative to the control: negative (red), neutral (grey), and positive (beneficial, stimulatory, or compensatory; green).

Notably, mean effect-size estimates for comparable traits sometimes differed among syntheses, even where primary studies were likely to overlap. This variation suggests that conclusions about microplastic effects may be sensitive not only to biological context, but also to meta-analytical decisions such as trait grouping, inclusion criteria, effect-size metrics, and moderator structure. Thus, even in the best-studied vertebrate group, microplastic exposure does not generate a uniform signature of organismal damage. Instead, microplastic exposure produces a trait-specific profile of impairment, neutrality, and, in some contexts, apparent compensatory or stimulatory responses.

Beyond fish, evidence remains more fragmented across other vertebrates. In mammals, research is dominated by laboratory mice and rats used for human health risk assessment (M. Liu et al., 2023). Amphibian research has also expanded over the past decade, reflecting growing concern over the vulnerability of this globally threatened group (Rahman et al., 2024). In mammals and amphibians, effects have been reported on multiple functional traits categories, including physiology and health, energy and metabolism, growth and development, behaviour and locomotion, and immunity, although evidence remains equivocal among studies (M. Liu et al., 2023; Mansfield et al., 2024). By contrast, evidence for birds and reptiles remains comparatively limited, with most studies documenting microplastic occurrence and ingestion rather than consequences on health and performance (Altunışık et al., 2024; Mansfield et al., 2024).

### 2.2 Invertebrates

Invertebrates provide the broadest meta-analytical evidence base, spanning both aquatic and terrestrial systems and encompassing 3,590 effect sizes across a wide range of biological traits. Available syntheses include 95 aquatic and 25 terrestrial species, representing eight major taxa: Arthropoda, Mollusca, Cnidaria, Annelida, Echinodermata, Rotifera, Nematoda, and Tunicata (Abbas et al., 2025; Bao et al., 2025; Berlino et al., 2023; Funke et al., 2024; Ji et al., 2021; Li et al., 2024; Mason et al., 2022; Moyo, 2022).

In aquatic invertebrates, microplastics have a markedly negative mean effect on physiology and health (Figure 1H: Hedges g’ = -1.34 to -1.08). By contrast, the remaining response trait categories exhibit substantial variation, with mean effect sizes spanning negative, neutral, and, in some cases, positive values, including behaviour and locomotion (Figure 1A: Hedges g’ = -0.5 to -0.15), morphology (Figure 1G: Hedges g’ = -0.054), energy and metabolism (Figure 1B: Hedges g’ = -0.26 to 0.24), feeding and particle processing (Figure 1C: Hedges g’ = -1.34 to 2.68), growth and development (Figure 1D: Hedges g’ = - 2.52 to 0.98), reproduction (Figure 1I: Hedges g’ = -3.07 to 0.42; MD = -10.51 (Berlino et al., 2023)), and survival (Figure 1J: Hedges g’ = -1.26 to 0.5). This variability is evident both among trait categories and among taxa. Negative mean effects are particularly pronounced within specific taxa, including reduced growth in gastropods (Figure 1D: Hedges g’ = -1.26) and rotifers (Figure 1D: Hedges g’ = -2.52), and reduced reproduction in nematodes (Figure 1I: Hedges g’ = -3.07). Conversely, positive responses, although less frequent and often based on fewer estimates, are also reported in insect growth, feeding, and reproduction (Figure 1C,D,I: Hedges g’ = 0.98, 2.68, and 0.11, respectively), rotifer reproduction (Figure 1I: Hedges g’ = 0.42), and gastropod survival (Figure 1J: Hedges g’ = 0.5). Together, these findings indicate that strong negative effects are not universal or uniformly distributed across taxa and biological traits, with sensitivity varying substantially among invertebrate groups.

Terrestrial invertebrates show a comparable sequence of impairment and neutrality. Negative mean effects sizes are reported in behaviour and locomotion (Figure S2A: lnRR = - 0.36), feeding and particle processing (Figure 1C: Hedges’ g = -0.68; Figure S2B: lnRR = - 0.13), growth and development (Figure 1D: Hedges’ g = -0.7; Figure S2C: lnRR = -0.09), reproduction (Figure 1I: Hedges’ g = -3.28), physiology and health (Figure S2F: lnRR = - 0.31) and survival (Figure 1J: Hedges’ g = -1.17; Figure S2H: lnRR = -0.15; Figure S3E: OR = 0.49). Yet neutral responses again occur within the same evidence base, particularly for fecundity (Figure 1I: Hedges’ g = -0.46; Figure S2G: lnRR = -0.22), health (Figure 1H: Hedges’ g = 0.25), growth and development (Figure S2C: lnRR = - 0.07), and behaviour (Figure 1A: Hedges’ g = -0.24) in some insect syntheses.

Overall, the current meta-analytical evidence reveals substantial variation in animal responses to microplastic exposure. Rather than a consistent pattern of toxicity, responses range from clear impairment to weak or neutral outcomes and, in some cases, apparent stimulatory responses. Such variation suggests that the biological consequences of microplastic exposure are shaped by interactions among multiple, interacting factors rather than by exposure alone. Understanding the sources of this variation remains a major challenge. In the following sections, we synthesize how exposure conditions, particle characteristics, species biology, and physiological responses may contribute to these contrasting outcomes.

## 3. Sources of variation in microplastic effects

### 3.1 Particle properties and exposure regimes

Microplastic effects are thought to highly depend on the properties of the particles themselves and on the exposure context in which organisms encounter them. Current evidence shows that biological responses are shaped by microplastic dose, exposure duration, polymer type, particle size, shape, density, ageing state, and biological colonisation (Figure 2; Abbas et al., 2025; Bao et al., 2025; Salerno et al., 2021; Wang et al., 2024). These factors may determine not only how likely particles are to be encountered and ingested, but also whether they are retained, egested, translocated, or able to interact with tissues long enough to induce measurable responses.

**Figure 2.**
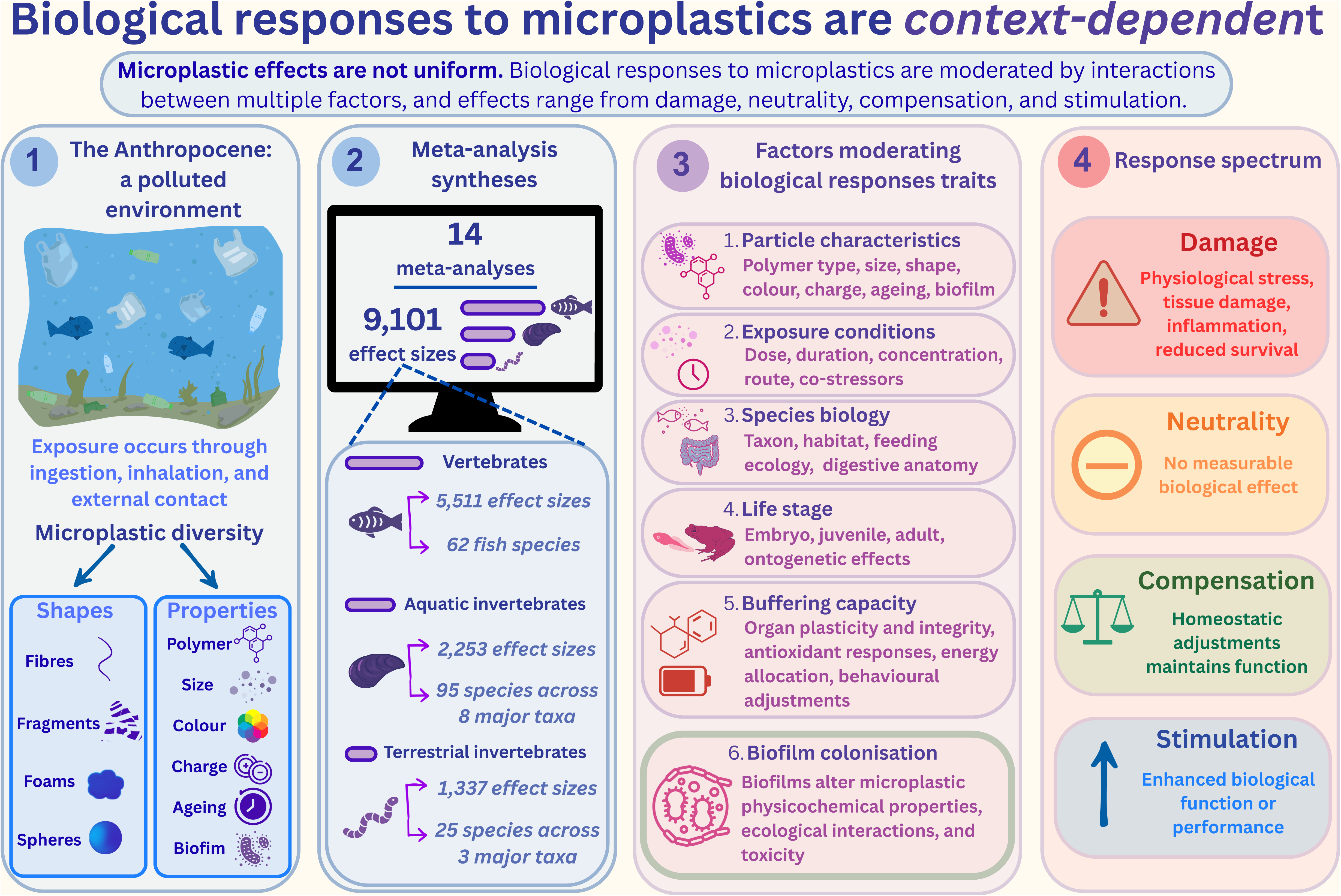
Conceptual framework summarising the context-dependent nature of biological responses to microplastic exposure. (1) Microplastics comprise heterogeneous particle mixtures that enter organisms through multiple exposure pathways. (2) Fourteen meta-analyses comprising 9,078 effect sizes demonstrate that responses vary across vertebrate and invertebrate taxa. (3) Biological outcomes are moderated by interacting factors including particle properties, exposure conditions, species biology, life stage, buffering capacity, and biofilm colonisation. (4) Responses span a continuum from damage through neutrality and compensation to stimulation.

#### 3.1.1 Particle characteristics

Polymer type, shape, size, and colour are key determinants of the biological impact of microplastics across vertebrates and invertebrates (Abbas et al., 2025; Bucci et al., 2020; Sacco et al., 2026; Thompson et al., 2024; Thornton Hampton et al., 2022). Among these characteristics, polymer type has been strongly linked to variation in life-history and health-related responses, with polyethylene and polystyrene displaying pronounced negative effects on growth and survival in fish, and health and development in aquatic and terrestrial invertebrates (Abbas et al., 2025; Bao et al., 2025; Sacco et al., 2026). Certain polymers such as polyethylene and polystyrene, are disproportionately represented in experimental approaches, meaning that apparent polymer-specific effects may partly reflect research effort rather than ecological prevalence or intrinsic toxicity (Hossain & Olden, 2022). This imbalance is important because microplastic risk assessments often rely on experimental evidence generated from a limited subset of particle types.

Microplastic size and density influence vertical distribution in aquatic systems and suspension in soils or air, thereby shaping which organisms encounter which particles (Klein et al., 2023; Porter et al., 2023; Stride et al., 2024; Yacoub & Han, 2026). Particle size is an important moderator of particle toxicity, and smaller particles are generally associated with higher toxicity (Bucci et al., 2020; Thornton Hampton et al., 2022). Smaller particles are more readily available to animals, including those with smaller body sizes, or smaller mouth openings (Jâms et al., 2020; Salerno et al., 2021). Their larger surface-area-to-volume ratio may also facilitate translocation across the gut epithelium and accumulation in secondary organs via the bloodstream (Cattaneo et al., 2023). Particles within the size range of 1-5 μm can cross epithelial barriers: in *Danio rerio*, microbeads can translocate across the intestinal epithelium and accumulate in the liver (Cattaneo et al., 2023). Beyond size, shape can further influence toxicity, and irregularly shaped particles, including fibres and fragments, may increase physical abrasion, reduce egestion efficiency, and prolong gut retention because of their surface structure and morphology (Mason et al., 2022). Fibres and fragments dominate environmental samples, and yet microspheres are disproportionately represented in laboratory experiments (Hossain & Olden, 2022). Colour is an additional, less explored particle characteristic that may shape ingestion risk and biological effects. Microplastics can be similar in colour to natural prey, potentially increasing the likelihood of ingestion (Niu et al., 2026; Sacco et al., 2026). In addition, the artificial dyes used to colour microplastics consist of organic and inorganic pigments that alter the properties and toxicity of the particles, and the original colour can change during environmental weathering (Niu et al., 2026; Sacco et al., 2026; Wei et al., 2025). Thus, microplastic exposure is not simply a matter of presence or absence, but of particle-specific encounter, uptake, retention, and biological reactivity.

#### 3.1.2 Dose and exposure duration

Higher microplastic concentrations are generally expected to produce stronger biological effects; however, the use of environmentally unrealistic concentrations and exposure durations remains a major limitation of the field (Bucci et al., 2020; Burns & Boxall, 2018; Cunningham & Sigwart, 2019). Experimental studies often represent worst-case or projected future scenarios rather than concentrations typically encountered in natural systems (Agathokleous et al., 2021). Conversely, short-term experiments may underestimate chronic effects, delayed ontogenetic costs, or non-linear responses such as compensation and hormesis, which may only emerge after chronic or repeated exposure (Agathokleous et al., 2021; Bucci et al., 2020; Ruthsatz et al., 2023). Thus, dose and duration jointly determine whether microplastic exposure is likely to remain biologically neutral, activate buffering responses, or induce damage.

### 3.2 Species traits shaping exposure and sensitivity

Plastic particles must first be encountered, ingested or inhaled, retained long enough to interact with tissues, and occur at doses capable of inducing measurable responses (Agathokleous et al., 2021; Bucci et al., 2020). For instance, aquatic vertebrates and invertebrates can experience continuous exposure through both water and prey ingestion, particularly in pelagic and benthic systems where filter-feeding increases encounter rates because microplastics can overlap in size and appearance with natural food particles (Hossain & Olden, 2022; Mason et al., 2022; Moyo, 2022; Porter et al., 2023). By contrast, many terrestrial organisms are likely exposed more episodically through diet, contaminated soil, or inhalation, although these routes may still be substantial in highly contaminated habitats (M. Liu et al., 2023; Mansfield et al., 2024). Habitat therefore shapes not only exposure intensity, but also whether exposure is continuous, pulsed, dietary, respiratory, or contact-based (Figure 2).

Ontogenetic changes in body size, mouthpart morphology, diet, habitat use, and gut structure can determine which particles are ingested and how efficiently they are cleared (Jâms et al., 2020; Salerno et al., 2021). For example, early larval mouthparts may be too small to ingest some particles, whereas larger mouthparts at later developmental stages may increase the size overlap between microplastics and natural prey, thereby raising ingestion likelihood (Jâms et al., 2020; Lievens et al., 2023). In fish, both the impact of microplastics and the size range of particles ingested depend on larval size and mouthpart morphology (Salerno et al., 2021). Life-history transitions may also reset or reduce exposure. In semi-aquatic species, aquatic larvae may experience sustained exposure, whereas metamorphosis and movement into terrestrial habitats could reduce microplastic loads through tissue remodelling, gut clearance, or shifts in habitat and diet (Szkudlarek et al., 2023).

Microplastic exposure therefore varies systematically with habitat, feeding ecology, particle properties, anatomy, and life-history stage, shaping encounter rates, retention time, and the probability of tissue interaction (Figure 2). Apparent taxonomic differences in microplastic sensitivity may partly reflect differences in exposure, uptake, and retention rather than intrinsic physiological vulnerability alone. These sources of variation provide the context for the next section, which examines how organisms may actively buffer microplastic exposure once particles have been encountered or internalised.

## 4. Buffering responses to microplastic exposure

### 4.1 Digestive buffering mechanisms

The digestive tract is the main interface between animals and ingested microplastics: particle clearance, gut plasticity, and barrier integrity (Jovanović et al., 2018; Ruthsatz et al., 2022; Vancamelbeke & Vermeire, 2017). Once particles enter the gut, outcomes depend on whether they are rapidly egested, retained long enough to affect nutrient acquisition, or brought into sustained contact with the intestinal lining (Kaur et al., 2024). These digestive filters and buffering mechanisms may primarily explain why microplastic ingestion sometimes causes weak, neutral, or even stimulatory responses (Figure 2).

#### 4.1.1 Gut clearance

Microplastics often accumulate in the gut which can reduce nutrient availability and affect energetically demanding processes such as growth and reproduction (Abbas et al., 2025; Berlino et al., 2023; Moyo, 2022; Salerno et al., 2021; Wang et al., 2024). In this context, rapid gut clearance can limit effective exposure by shortening retention time. In filter-feeding tadpoles of *Xenopus tropicalis*, most ingested microplastics were egested within six hours and almost completely cleared after six days (L. Hu et al., 2016). Similarly, in *Sparus aurata*, gut retention was low after 45 days of exposure to six microplastic types followed by 30 days of depuration (Jovanović et al., 2018). Mechanical processing may also affect particle fate. In birds, gizzards routinely process indigestible material, and microplastics can be fragmented and subsequently egested, although whether fragmentation facilitates clearance or instead generates smaller, more bioavailable particles remains unresolved (Ramos-Elvira et al., 2025; Terepocki et al., 2017; Winkler et al., 2020). Comparable structures, such as earthworm gizzards or crustacean gastric mills, can also influence particle breakdown and egestion, but their buffering role remains poorly tested (LeBlanc et al., 2025; Meng et al., 2023).

#### 4.1.2 Gut plasticity

Optimal digestion theory predicts that, under nutrient-dilute conditions, organisms may increase gut length and prolong retention time to maximise nutrient extraction efficiency (Sibly, 1981). By adding non-nutritive dietary material to the gut contents, ingested microplastics can elicit compensatory digestive responses (Ruthsatz et al., 2022). In *Xenopus laevis* larvae, exposure to pristine microplastics increased gut length and gut mass, a response also observed in larvae fed cellulose, an indigestible plant fibre. In this example, larval body condition was maintained despite reduced body mass, suggesting that gut enlargement may have partially compensated for energetic intake by increasing digestive capacity (Ruthsatz et al., 2022). Likewise, juvenile fish exposed to microplastics also showed increased densities of mucosal goblet cells at higher concentrations, suggesting plastic reinforcement of the intestinal lining (Pedà et al., 2016). Such plastic responses could help to mitigate short-term effects, but they could also alter particle residence time or impose maintenance costs that emerge later in development. Whether these responses improve clearance, nutrient uptake, or long-term protection from processes such as abrasion remains unresolved.

#### 4.1.3 Barrier integrity

Barrier integrity determines whether microplastic effects remain local or progress towards tissue damage and systemic exposure. The intestinal epithelium regulates selective permeability, whereas mucus-secreting goblet cells form a protective layer that can reduce direct contact between particles and epithelial cells and facilitate particle transport through the gut (Liquori et al., 2007; Vancamelbeke & Vermeire, 2017). Together, these features form an intrinsic buffering mechanism that limits particle penetration and tissue damage. In some species, barrier function may also be reinforced through plastic responses, such as increased goblet cell density or mucus production (Pedà et al., 2016). When particles are retained, sustained contact with the intestinal lining may promote abrasion, inflammation, increased permeability, or translocation across the barrier (Kaur et al., 2024). Yet these responses are likely to differ among species depending on their natural exposure to suspended particles, gut structure, and epithelial defences. In *Oncorhynchus mykiss*, a species inhabiting environments with high turbidity and chronic exposure to suspended inorganic particles, exposure to relatively large polystyrene microplastics for four weeks did not alter intestinal permeability, active transport, or electrophysiological parameters (Ašmonaitė et al., 2018). The ability to maintain intestinal functioning under microplastic exposure may reflect the capacity to tolerate particulate exposure, potentially associated with their adaptation to naturally particle-rich, turbid environments. Similarly, chronic low-dose exposure to polyethylene terephthalate microplastics in mice altered immune-associated transcriptional profiles without detectable changes in intestinal pathology or mucin barrier integrity (Harusato et al., 2023). Together, these findings suggest that physiological or molecular responses to microplastics do not necessarily lead to the disruption of the intestinal barrier function, and existing barrier defences may buffer against tissue-level damage under certain conditions. Gut microbiota and probiotics may further modulate microplastic-gut interactions, although these pathways remain poorly understood (Fackelmann et al., 2023; Gao et al., 2025; Teng et al., 2025).

### 4.2 Antioxidant buffering responses

At the cellular level, microplastic exposure can promote reactive oxygen species production through several interconnected pathways, including particle-tissue interactions, mitochondrial dysfunction, immune activation, and microbiome-mediated processes (M. Hu & Palić, 2020). When reactive oxygen species exceed antioxidant capacity, they can cause damage to biomolecules including proteins, lipids, and DNA (Halliwell & Gutteridge, 2015; Pisoschi & Pop, 2015). However, antioxidant systems can buffer these effects, with enzymes such as superoxide dismutase and catalase acting synergistically to maintain redox homeostasis and prevent oxidative damage when exposure remains within compensatory limits (Ighodaro & Akinloye, 2018; Mates, 1999; Pamplona & Costantini, 2011).

Across vertebrates and invertebrates, microplastic studies report three broad oxidative-response profiles, including no detectable change in oxidative biomarkers, increased antioxidant activity without measurable oxidative damage, and elevated oxidative damage under high dose or chronic exposure regimes (Bao et al., 2025; Hossain & Olden, 2022; Salerno et al., 2021; Wang et al., 2024). These patterns suggest a continuum from no detectable oxidative response, through antioxidant activation consistent with successful buffering, to oxidative damage when compensatory capacity is exceeded. For example, environmentally realistic microplastic exposure can induce antioxidant responses without detectable oxidative damage in juvenile frogs, suggesting effective antioxidant buffering responses (Park et al., 2024). In contrast, evidence indicates that oxidative damage can increase under some exposure contexts, particularly when exposure intensity, duration, particle properties, or organismal vulnerability exceed compensatory capacity (Bao et al., 2025; Hossain & Olden, 2022; Salerno et al., 2021; Wang et al., 2024). This balance between activation and damage may help explain why some studies report neutral organismal outcomes despite clear molecular or biochemical responses (Figure 2).

Antioxidant buffering is also linked to broader cellular maintenance pathways such as immune regulation, mitochondrial function, proteostasis, DNA repair, in addition to ageing-related mechanisms such as telomere dynamics (Halliwell & Gutteridge, 2015). Furthermore, since buffering oxidative damage requires energy and resources that could otherwise sustain development or reproduction (Burraco et al., 2022; Halliwell & Gutteridge, 2015; Metcalfe & Monaghan, 2001), microplastic exposure may generate fitness-related costs. Conversely, mild activation of antioxidant and cellular repair pathways could produce hormetic-like (low-dose stimulation) outcomes if it enhances stress resistance without imposing substantial costs (Martin et al., 2024, 2025). Whether such responses represent adaptive buffering and short-term compensation, or early signs of future damage will depend on exposure duration, life stage, tissue type, and the energetic state of the organism. Delayed consequences may be particularly likely where exposure persists across life stages. Microplastics can persist across developmental transitions through ontogenetic transfer, extending exposure beyond the initial life stage and potentially increasing the likelihood of delayed or cumulative effects (Alizadeh et al., 2025; Ruthsatz et al., 2023). Future work should determine whether, and to what extent, redox regulation enables organisms to maintain health and fitness under microplastic exposure.

### 4.3 Behavioural buffering responses

Behavioural changes under microplastic exposure may represent adaptive responses that reduce particle uptake or mitigate the energetic and physiological consequences of exposure (Figure 2). Microplastics can resemble the size or chemical cues of natural prey items, leading some organisms to mistakenly ingest them (Fabra et al., 2021; Hamann et al., 2024; Vroom et al., 2017). However, shifts in foraging behaviour may influence microplastic exposure through selective feeding and particle discrimination. Some aquatic invertebrates, including rotifers (Bermúdez et al., 2025), copepods (Traboni et al., 2023), amphipods (Yardy & Callaghan, 2020), and corals (Joppien et al., 2022), can differentiate between and preferentially select food particles over microplastics under favourable feeding conditions. Similar particle discrimination has also been observed in fish, including *Oryzias melastigma* (Lan et al., 2026), and such feeding selectivity may reduce microplastic ingestion. However, its effectiveness is often reduced under low-food conditions, when nutritional demands increase the likelihood of microplastic ingestion (Lan et al., 2026), under high particle concentrations (Bermúdez et al., 2025), or when organisms are exposed to aged particles (Joppien et al., 2022). Evidence for similar behavioural buffering responses across taxa remains limited, particularly in terrestrial ecosystems.

## 5. Biofilm colonisation as a mediator of microplastic effects

Microplastics are rarely encountered as pristine particles in natural environments. Once released, their surfaces are modified by physical, chemical, and biological processes, which commonly include biofilm colonisation (Figure 3; P. Liu et al., 2020; Moyal et al., 2023). Biofilms can alter microplastic physicochemical properties, influencing particle transport, the probability of ingestion, and how microplastics interact with animal tissues (Figure 2; Fabra et al., 2021; Hamann et al., 2024; Vroom et al., 2017).

**Figure 3.**
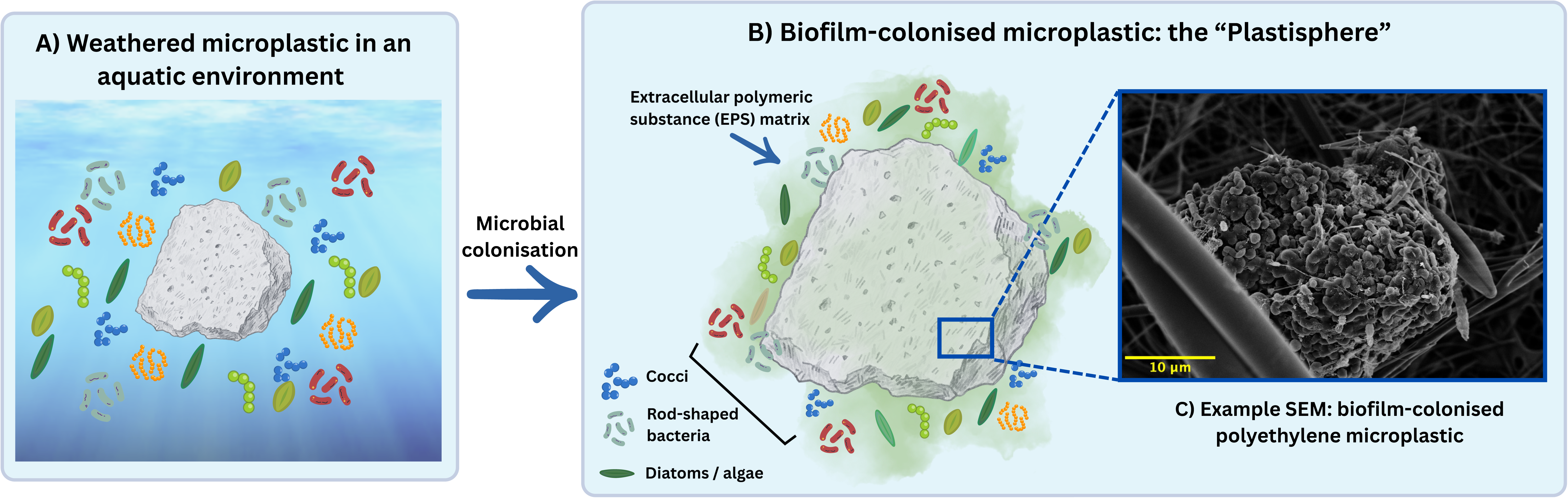
Biofilm formation on microplastics in aquatic environments. A) Weathered microplastic particles in aquatic environments are surrounded by free-living microorganisms, including fungi, bacteria, algae, and protists, that can colonise the particle surface. B) Following microbial colonisation, a structured biofilm community develops on the microplastic surface, comprising cocci, rod-shaped bacteria, diatoms, and algae embedded within an extracellular polymeric substance (EPS) matrix that supports and stabilises the biofilm. C) Example of a scanning electron microscope (SEM) image showing microbial colonisation of a pristine polyethylene microplastic following one month of ageing in pond water.

Biofilms forming on microplastic surfaces can comprise bacteria, archaea, fungi, and protozoa embedded within an extracellular matrix, creating an ecological niche known as the ‘plastisphere’ (Figure 3; Zettler et al., 2013). This biological coating can alter particle buoyancy, surface charge, sorption capacity, likelihood of ingestion, and the potential of microplastics to act as vectors for chemical pollutants or pathogens (P. Liu et al., 2020; Moyal et al., 2023; Wang et al., 2021). In this context, aged microplastics refer to particles whose surfaces have been modified by environmental processes such as UV exposure, mechanical abrasion, chemical weathering, wastewater incubation, or microbial colonisation, making them more representative of particles encountered in natural systems than pristine laboratory plastics (Rubin et al., 2021; Shi et al., 2024). For example, microplastics experimentally aged in water have been shown to attenuate multigenerational effects in *Daphnia magna* compared with pristine microplastics, suggesting that particle condition can alter toxicity and modify biological responses (Schür et al., 2021). Another study on the same species found that biofilm-colonised microplastics altered mortality relative to pristine particles (Motiei et al., 2021). In some vertebrate species, biofilm-colonised microplastics appear to modify physiological responses. For instance, in *Seriola lalandi,* pristine particles reduced aerobic scope and increased oxidative stress compared to biofilm-colonised particles, suggesting greater physiological costs of exposure (Kelly et al., 2024). The reduced fitness costs associated with biofilm-colonised particles may reflect the nutritional value of the biofilm-associated microorganisms, which could partially offset the energetic costs of ingesting pristine, non-nutritive particles. Although still limited, current evidence suggests that particle ageing and biological colonisation can modify the direction and magnitude of microplastic effects.

Biofilms may also change exposure by making particles more food-like. Biofilm-colonised microplastics can release chemical cues resembling those associated with natural food sources, thereby increasing ingestion likelihood (Fabra et al., 2021; Hamann et al., 2024; Vroom et al., 2017). Also, biofilms may alter the nutritional profile of ingested particles, potentially changing microplastics from purely indigestible diluents into particles carrying microbial biomass or associated nutrients (Horváthová & Bauchinger, 2019). This is particularly relevant under low-food conditions, where the energetic consequences of ingesting microplastics may depend on whether particles are pristine, aged, or biologically colonised. Biofilm has been shown to mitigate the nutritional dilution caused by microplastics in *Daphnia magna*, resulting in enhanced growth and survival compared with pristine plastic particles under low-food conditions (Amariei et al., 2022). This does not necessarily imply that biofilm-colonised microplastics are beneficial; rather, it suggests that microbial colonisation can change the sign and magnitude of microplastic effects depending on food availability, microbial community composition, particle properties, and consumer digestive physiology.

Biofilm colonisation therefore adds a missing ecological layer to microplastic research. It links particle ageing, microbial ecology, feeding behaviour, nutritional ecology, and toxicology, and may help explain why pristine microplastics used in laboratory experiments sometimes produce responses that differ from those observed under more realistic environmental conditions. Future studies should explicitly compare pristine, aged, and biofilm-colonised particles across food regimes, taxa, and life stages, while characterising biofilm composition and function. Doing so will be essential for understanding whether biofilms amplify toxicity, buffer nutritional costs, alter ingestion risk, or generate context-dependent responses in polluted ecosystems.

## 6. Future directions and conclusion

Microplastic research has largely emphasised negative effects on animal health, physiology, growth, reproduction, survival, and behaviour. Our synthesis of meta-analyses instead reveals a heterogeneous response landscape, with evidence for harm but also recurrent neutral and compensatory or stimulatory responses across taxa, traits, and ecological contexts (Figure 2). While neutral outcomes may be underrepresented because of publication bias, their recurrence across independent syntheses suggests that neutrality is not simply a statistical artefact, but a biological pattern requiring explanation. In addition, our study highlights that biological responses to microplastics are shaped by particle characteristics, physicochemical properties, and exposure regimes. Species-specific traits further influence exposure, uptake, retention, and the capacity for morphological and physiological buffering. The key question is therefore no longer whether microplastics are harmful, but when, why, and for whom they become harmful.

Future research should ideally prioritise more realistic and mechanistic experimental designs. Many studies still use pristine microspheres, whereas organisms in nature more often encounter fibres, irregular fragments, mixed polymer types, and aged biofilm-colonised plastics. Exposure concentrations also often exceed environmentally relevant levels, approximating worst-case rather than typical scenarios (Bucci et al., 2020; Hossain & Olden, 2022; Rahman et al., 2024). Experiments could therefore compare pristine, aged, and biofilm-colonised particles, single- and mixed-particle exposures, and concentrations that span current, future, and worst-case scenarios. Another priority is to determine how long buffering responses can be maintained under chronic and environmentally realistic exposure, and whether compensatory responses or their associated costs give rise to delayed effects across ontogeny. Short-term experiments may detect immediate responses but rarely show whether neutral outcomes reflect genuine tolerance, temporary compensation, or delayed ontogenetic costs. Longitudinal (i.e., within-individual) designs are needed to link early mechanisms such as gut plasticity, antioxidant activation, immune modulation, behavioural avoidance, or altered feeding to later outcomes in growth, development, reproduction, survival, ageing, and lifespan. Manipulating candidate buffering pathways can determine whether they mediate tolerance or compensation under microplastic exposure. Where possible, such designs should be extended across generations to test whether microplastic exposure produces transgenerational effects on offspring performance and resilience. Finally, because microplastics often occur alongside other global change stressors, multi-factor experiments will be needed to test whether buffering capacity is maintained under warming, food limitation, pathogens, contaminants, hypoxia, or other global-change stressors. These approaches should move beyond single endpoints towards multi-level assessments that could integrate measures of particle uptake and retention, gut morphology and barrier integrity, antioxidant and immune regulation, microbiome composition, behaviour, energy allocation, reproduction, survival, and other health-related markers. Such designs will help distinguish true absence of effect, successful compensation, hormetic-like stimulation, and early warning signs of delayed vulnerability.

Microplastics should be understood as context-dependent ecological and physiological challenges rather than uniformly toxic or biologically inert particles. Their effects emerge from interactions among particle properties, exposure realism, species biology, and buffering capacity. Recognising this complexity makes risk assessment more precise by identifying the conditions under which exposure produces damage, neutrality, compensation, or stimulation in polluted ecosystems.

## Funding

The German Research Foundation (DFG) project (No. 459850971) *A new actor on the stage of global change: A multi-level perspective on the toxicity of microplastics pollution in amphibians* supported CM and the research conducted here. KR was supported by a Marie Curie Fellowship AMPHISTRESS-101151070. PB was supported by Ramón y Cajal fellowship 2023-044964-I (Spanish Ministry of Science and Innovation).

## Competing interests

The authors have no competing interest to declare.

## Data availability statement

https://doi.org/10.6084/m9.figshare.33112100

## Author contributions

**CM**: Conceptualization, Methodology, Investigation, Data Curation, Formal Analysis, Visualization, Writing – Original Draft.

**PB**: Conceptualization, Methodology, Formal Analysis, Supervision, Writing – Review & Editing.

**KR**: Conceptualization, Methodology, Formal Analysis, Supervision, Funding Acquisition, Writing – Review & Editing.

## Notes

https://doi.org/10.6084/m9.figshare.33112100

### Competing Interest Statement

The authors have declared no competing interest.

