## Supplementary Material for "Beyond harm: synthesising meta-analytical and mechanistic evidence on animal responses to microplastics"

**Identification of studies via databases**

Records removed *before screening*:

Duplicate records removed (n = 101 )

Records identified from databases (n = 202)

**Identification**

Records screened

(n = 101)

Records excluded (n = 32)

Full-text articles excluded (n = 55):

1. Microplastics not evaluated as a single stressor
2. No quantitative effect-size measure reported
3. No relevant biological response reported

**Screening**

Studies included in review

(n = 14)

Reports assessed for eligibility

(n = 69)

**Included**

**Figure S1:** PRISMA flow diagram outlining the article selection process.

**Table S1:** Classification of biological response traits extracted from meta-analyses into major response categories used throughout the synthesis.

| **Final Category** | **Sub-category** | **Animal type** | **Meta-analysis reference** | **Biological response trait definitions** |
| --- | --- | --- | --- | --- |
| **Behaviour and locomotion** | Behaviour | Fish | Salerno et al. 2021 | Activity rate; locomotor activity; mobility; swimming velocity; maximum velocity; distance travelled; spontaneous movement; turning behaviour; inactivity. |
|  |  | Fish | Hossain & Olden 2022 | Foraging (feeding time, foraging time, and number of bites); mobility (distance travelled, mean speed, dark phase/photo phase, and resting individuals). |
|  |  | Fish and aquatic invertebrates | Berlino et al. 2023 | Activity rate; locomotor activity; mobility; swimming velocity; maximum velocity; distance travelled; spontaneous movement; turning behaviour; inactivity; use of the tank; predatory activity; feeding success; foraging time. |
|  |  | Aquatic invertebrates | Mason et al. 2022 | Righting time; byssal thread production; cirral beating frequency; swimming speed. |
|  |  | Terrestrial invertebrates | Abbas et al. 2025 | Case length; climbing activity; food selection; food selection rate; interior width; locomotory activity; residence time; sleep time; sum activity. |
|  |  | Terrestrial invertebrates | Li et al. 2024 | Anterior width; case length; case weight; climbing ability; curling rate; food choice; food selection rate; locomotor activity; neuromuscular junction alteration; residence time; retinal response; sleep time; sum activity. |
|  | Behavioural, sensory and neuromuscular functions | Fish | Yuan et al. 2023 | Predatory performance; swimming speed; serotonin (5-HT); acetylcholinesterase (AChE) activity; eye and photoreception development-related gene expression (six3, pax6, rx3, rh1, lws, and sws1); nervous system development-related gene expression (α1-tubulin, elavl3, gap43, gfap, mbp, syn2a, lfng, and olig2). |
|  |  | Fish and aquatic invertebrates | Bao et al. 2025 | Thoracic limb activity; swimming speed; swimming distance; acetylcholinesterase (AChE) activity; time spent in the social segment; time spent not moving; filtration rate; ingestion rate. |
|  | Locomotion | Fish | Yuan et al. 2025 | Absolute turn angle; active time; anxiety index; body contact; burst swimming; cluster scores; distance during feeding time; distance to the centre of the well; exploration; fin flickering; hatching rate; hatching time; instantaneous speed; jerking; larvae resting on their side; larval density; latency to enter the top; manic time; mean absolute turn angle; mean activity; mean distance between fish; mobility; moved distance; movement; movement delay; movement duration; moving time; no movement; percentage of time that fish remained in the centre zone; percentage of time that fish remained in the dark zone; prey–predator distance; shoal area; stasis; static time; swimming area; swimming speed; percentage of time that larvae spent in the upper area of the well with stimulus; percentage of time that larvae spent in the upper area of the well without stimulus; thigmotaxis; time spent close to the shelter; time spent far from the shelter; time spent in the shelter; time spent in the top; total crossings; velocity. |
|  | Mobility | Fish | Sacco et al. 2026 | Swimming velocity; distance travelled. |
| **Energy and metabolism** | Energy consumption | Aquatic invertebrates | Mason et al. 2022 | Respiration rate (oxygen consumption); energy consumption. |
|  | Metabolism | Fish | Yuan et al. 2023 | 7-Ethoxyresorufin O-deethylase activity (EROD); 7-benzyloxy-4-trifluoromethyl-coumarin O-dibenzyloxylase activity (BFCOD); metallothionein; amino acid metabolism; carbohydrate metabolism; lipid metabolism; glycogen content; protein content; P-glycoprotein content; cyp1a (detoxification); glucose; pyruvate; glycolipid metabolism-related gene expression (Gk, Hk1, Aco, PPar-α, Cpt1, Acc, Fas, PPar-γ, and Apo); total cholesterol; triglyceride content; energy-related enzyme activity (lactate dehydrogenase and isocitrate dehydrogenase). |
|  |  | Fish | Yuan et al. 2025 | Gonadosomatic index; heart rate; heartbeats; hepatosomatic index; respiratory rate; visceral somatic index; viscerosomatic index. |
|  |  | Fish and aquatic invertebrates | Bao et al. 2025 | Na⁺/K⁺-ATPase activity; Ca²⁺/Mg²⁺-ATPase activity; alginose content; non-esterified fatty acid content; triglyceride content; cholesterol content; bile acid content; caspase-3 activity; caspase-9 activity; protein content; soluble sugar content; glutamic acid content; asparagine content; arginine content; 7-ethoxyresorufin O-deethylase activity; 7-benzyloxy-4-trifluoromethyl-coumarin O-dibenzyloxylase activity; maximum quantum yield of photosystem II (PSII) (Fv/Fm); electron transport rate; metabolism-related gene expression (cyp314, hr96, p-gp, mt, cyp, lys, p53, act, cel.1, cel.2, abcc2, soat2, cyp7a1, cyp8b1, dgat2, lpl, mttp, apoba, apobb1, apoa1a, apoa4b2, slc25a25a, casp3, casp9, baxa, cyp1a, cyp24a1, cyp2p8, abcc3, abcc5, mfn2, defbl1 and abcb4). |
|  |  | Fish and aquatic invertebrates | Berlino et al. 2023 | Energy consumption; oxygen consumption; assimilation efficiency; macronutrient content; condition factor; hepatosomatic index; gonadosomatic index; heart rate; mucus secretion; byssus secretion |
|  | Nutrition | Fish | Yuan et al. 2025 | Condition factor |
| **Feeding and ingestion** | Consumption | Fish | Hossain et al. 2022 | Predatory performance (percentage of supplied prey ingested). |
|  | Feeding | Fish | Sacco et al. 2026 | Food acquisition behaviours: searching, capturing, handling, and consuming prey. |
|  |  | Fish | Salerno et al. 2021 | Predatory performance; feeding success; foraging time; number of ingested prey. |
|  |  | Fish | Yuan et al. 2025 | Daily food intake; feed intake; feeding ability; feeding delay; feeding success; foraging time; ingested prey; number of bites; predatory performance; time to consume the given food. |
|  |  | Fish and aquatic invertebrates | Moyo 2022 | Foraging (selectivity, giving-up time, feeding time); ingestion rate; egestion (defaecation) rate; predation (% attacked, % caught, % consumed, number eaten). |
|  |  | Aquatic invertebrates | Mason et al. 2022 | Prey consumption rate; algal clearance rate; ingestion success (%). |
|  |  | Terrestrial invertebrates | Abbas et al. 2025 | Food intake. |
|  |  | Terrestrial invertebrates | Li et al. 2024 | Defecation rate; food intake. |
|  | MPs ingestion and accumulation | Fish and aquatic invertebrates | Bao et al 2025 | No description provided. |
| **Growth and development** | Development | Fish | Salerno et al. 2021 | Biometric measurements; head length; head height; head depth; liver weight; gill weight; gonad weight; swim bladder area; optic vesicle area; pericardium area; angle between myosepta; distance between myosepta; interocular distance. |
|  |  | Aquatic invertebrates | Mason et al. 2022 | Normal development (%); larval abnormalities (%); development time; segment regeneration time. |
|  |  | Terrestrial invertebrates | Abbas et al. 2025 | Adult emerged; development time; egg to pupae; embryo to eclosure; emerged individual number; emergence ratio; emerging time; larval cycle; pupae emerged; stage duration. |
|  |  | Terrestrial invertebrates | Li et al. 2024 | Day to emergence; development time; egg to adult; egg to eclosure; egg to pupae; emerged individual number; emergence ratio; emergence time; larval cycle rate; pre-pupae stage rate; pupae length; stage duration. |
|  | Growth | Fish | Hossain et al. 2022 | Body weight; body length; condition factor; hepatosomatic index; growth rate. |
|  |  | Fish | Sacco et al. 2026 | Included weight- and length-based measurements. |
|  |  | Fish | Salerno et al. 2021 | Body weight; body length; standard length; total length; weight gain rate; body mass; changes in body mass; length–weight ratio. |
|  |  | Fish | Wang et al. 2024 | Body length; body weight; specific growth rate. |
|  |  | Fish | Yuan et al. 2025 | Absolute growth rate; body area; body depth; body length; body weight; body width; condition factor; embryo diameter; food conversion rate; heartbeats; muscle weight; specific growth rate; standard length; total length; total weight; weight gain; weight gain rate. |
|  |  | Fish and aquatic invertebrates | Berlino et al. 2023 | Body weight; body length; standard length; total length; weight gain rate; body mass; changes in body mass; length–weight ratio; biometric measurements; head length; head height; head depth; liver weight; gill weight; gonad weight; swim bladder area; optic vesicle area; pericardium area; angle between myosepta; distance between myosepta; interocular distance. |
|  |  | Fish and aquatic invertebrates | Moyo 2022 | Body condition; body size; growth rate; body length. |
|  |  | Aquatic invertebrates | Mason et al. 2022 | Somatic growth rate; length increase; weight increase. |
|  |  | Terrestrial invertebrates | Abbas et al. 2025 | Body length; body weight; frame cover (adult); frame number used; growth rate; larval cycle; left wing length; right wing length; wing length; wing size. |
|  |  | Terrestrial invertebrates | Ji et al. 2021 | Change in weight. |
|  |  | Terrestrial invertebrates | Li et al. 2024 | Body length; body weight; cocoon weight; frame with insect; frames used; growth rate; pupae weight; wing length. |
|  | Growth & development | Fish and aquatic invertebrates | Bao et al. 2025 | Survival rate; growth inhibition; body length; body weight; root length; biomass productivity; dry cell weight; malformation rate; heart rate; chlorophyll a content; chlorophyll b content; development-related gene expression (jhe, ecrb, col1a2, col11a2, col11a1a, col2a1b, cdx4, tnnc2, and SmyD1). |
|  |  | Fish | Yuan et al. 2023 | Length; weight; survival rate; death rate; deformity rate; hatching time; heartbeats; triiodothyronine; thyroxine; development-related gene expression (BMP4, NKX2.5, COX, EPO, FGF8, ATPase, GATA4, and SmyD1). |
| **Immunity** | Immunity | Fish | Hossain et al. 2022 | Alkaline phosphatase (ALP) activity; neutrophil degranulation (percentage increase); blood globulin concentration; leucocyte viability. |
|  | Immune system | Fish | Yuan et al. 2023 | Lysozyme activity; immunoglobulin M (IgM) level; nitric oxide (NO) content; inducible nitric oxide synthase (iNOS) protein; total immunoglobulins; alternative complement activity; acid phosphatase activity; alkaline phosphatase activity; complement 3 (C3); number of melanomacrophage centres; caspase-3 activity; D-lactate content; number of goblet cells (skin); inflammatory response-related gene expression (IL-1β, IL-6, TNF-α, tnfα, cc-chem, nf-kb, cox1, IFN, TGFβ1, MMP2, and HSP70). |
|  |  | Fish and aquatic invertebrates | Bao et al. 2025 | D-lactate content; lysozyme activity; immunoglobulin M (IgM) level; complement 3 (C3); inflammatory response-related gene expression (IL-1β, IL-6, TNF-α, tnf-α, NF-κB, NF-KB1, c-Rel, IFN, TNF-β, TGFβ1, TLR-2, TLR-4, MyD88, and cebpb). |
| **Microbiome** | Gut microbiome diversity | Fish | Wang et al 2024 | Gut microbiota diversity. |
| **Morphology** | Alimentary and excretory systems | Fish | Yuan et al. 2023 | Hepatic index; trypsin activity; amylase activity; lipase activity; aspartate aminotransferase activity; alanine aminotransferase activity; gamma-glutamyl transferase activity; alkaline phosphatase activity; lactate dehydrogenase activity; mucus coverage ratio; gill lamella height; intestinal bulb tunica muscularis thickness; intestinal bulb fold height. |
|  |  | Fish and aquatic invertebrates | Bao et al. 2025 | Mucus coverage rate; lipase activity; amylase activity; trypsin activity; hepatosomatic index; stomach fullness index; diamine oxidase activity; intestinal damage; intestinal villus length; number of goblet cells; intestinal microbiota-related indicators (Chao index, Shannon diversity); alimentary and excretory-related gene expression (CLDN5, Oclna, and ZO-1). |
|  | Exterior morphology | Fish | Yuan et al. 2025 | Eye size; head depth; head height; head length; head-to-trunk angle; head width; head-to-body length ratio; interocular distance; malformation rate; mouth distance; peduncle depth; caudal fin (%); abnormal fish in the caudal fin (%); normal fish in the caudal fin (%); proboscis length; relative area of the regenerating fin; tail length. |
|  | Interior morphology | Fish | Yuan et al. 2025 | Brain fraction (% of total fish); caudal fin complex deformity; caudal fin vertebral deformity; caudal vertebral deformity; oedema; intestinal enterocyte height; ventral epidermis; dorsal epidermis; intestinal fold height; gyrus size; intestine organ index; jaw organ index; liver organ index; malformation rate; intestinal microvillus height; morphological abnormality rate; axial skeleton deformities (%); abnormal fish with axial skeleton deformities (%); normal fish with axial skeleton deformities (%); pericardial oedema area; precaudal vertebral deformity; spine length; swim bladder area; brain water content; Weberian vertebral deformity; weight loss. |
|  | Intestines | Fish | Hossain et al. 2022 | Diameter/thickness; goblet cell concentration (cells/mm²; periodic acid–Schiff (PAS)-positive cells); gastrointestinal histopathology score (semi-quantitative scoring); mucus secretion coverage ratio (ratio of the mucus secretion area to the total colon area (%)); neutrophil concentration (cells/mm²); toll-like receptor 2 concentration; trypsin activity. |
| **Physiology & Health** | Health | Fish | Salerno et al. 2021 | Condition factor; hepatosomatic index; gonadosomatic index; heartbeat; mucus secretion; oxygen consumption. |
|  |  | Terrestrial invertebrates | Abbas et al. 2025 | Abaecin; ACE index; adrenic acid; antioxidant activity; apidaecin; arachidic acid; arachidonic acid; attacin; behenic acid; brood area; butyric acid; C20:2 fatty acid; caprylic acid; catalase (CAT); cecropin A; Chao index; Chao 1 index; Chao 2 index; Chao 3 index; Cyp Q1; Cyp Q2; D-glucose content; D-lactate (D-Lac); diamine oxidase (DAO); defensin; docosadienoic acid; docosahexaenoic acid; domeless; eicosapentaenoic acid; elaidic acid; γ-linolenic acid; gondoic acid; glutathione (GSH); glutathione S-transferase (GST); GstD1; GstD3; honey area; hopscotch; lauric acid; linoleic acid; linolenic acid; lipid peroxidation (LPO); lysozyme; margaric acid; malondialdehyde (MDA); myristic acid; myristoleic acid; nervonic acid; oleic acid; palmitic acid; palmitoleic acid; PD; pentadecylic acid; propionic acid; protein content; protein level, lipid level, and microelement level; reactive oxygen species (ROS); Shannon diversity; Shannon index; Simpson diversity; superoxide dismutase (SOD); stearic acid; symplekin; triglyceride (TG) content; vaccenic acid. |
|  |  | Terrestrial invertebrates | Li et al. 2024 | Bacterial microbiome; brood area; carbohydrate level; Chao 1 index; detoxification response; energy consumption; fungal mycobiome; general stress; honey area; immunisation stress; internal organ stress; lipid level; microelement level; oxidative stress; physical stress; protein level; repair gene expression; richness; Shannon index; uric acid; wing recombination. |
|  | Oxidative damage | Fish | Wang et al. 2024 | Malondialdehyde (MDA) level. |
|  |  | Fish | Yuan et al. 2023 | Lipid peroxidation; malondialdehyde (MDA) content; reactive oxygen species (ROS); protein carbonyl content; 8-hydroxy-2′-deoxyguanosine (8-OHdG) level; hydroperoxide content. |
|  |  | Fish and aquatic invertebrates | Bao et al. 2025 | Superoxide dismutase (SOD) activity; malondialdehyde (MDA) content; catalase (CAT) activity; reduced glutathione (GSH) content; glutathione S-transferase (GST) activity; reactive oxygen species (ROS); superoxide radical content; peroxidase activity; lactoperoxidase activity; hydroxyl radical content; total antioxidant capacity; oxidative stress-related gene expression (sod, cat, gpx1a, got2a, and gstp2). |
|  | Toxicity | Fish | Hossain et al. 2022 | Acetylcholinesterase (AChE) activity (concentration); catalase (CAT) level (concentration); oxidative burst (percentage increase); superoxide dismutase (SOD) activity (concentration). |
| **Reproduction** | Fecundity | Fish | Salerno et al. 2021 | Egg production; fertilisation rate. |
|  |  | Terrestrial invertebrates | Abbas et al. 2025 | Number of eggs; total offspring. |
|  |  | Terrestrial invertebrates | Li et al. 2024 | Egg number; total offspring. |
|  | Fertility | Fish | Sacco et al. 2026 | Number of eggs; proportion of fertile organisms. |
|  | Hatching | Fish | Salerno et al. 2021 | Hatching percentage; hatching success; hatching rate; hatching time. |
|  | Hatching rate | Fish | Sacco et al. 2026 | Percentage of hatched eggs. |
|  | Reproduction | Fish | Hossain & Olden 2022 | Fecundity (mean number of eggs per female); gonadosomatic index; hatching rate (time from fertilisation to hatching; hatching success (%)). |
|  |  | Fish | Wang et al. 2024 | Average number of eggs laid; hatching rate; hatching time. |
|  |  | Fish | Yuan et al. 2023 | 17β-estradiol; testosterone; vitellogenin; egg number per female per day; gonadosomatic index; mature sperm count; atretic follicles (%); offspring indicators (growth and development); reproduction-related gene expression (VTG1, CHgH, ERα, Gthα, fshβ, lhβ, mGnrh, and 3βHSD). |
|  |  | Fish | Yuan et al. 2025 | Fertilisation rate; hatching time; hatching rate; number of eggs; number of hatched embryos; sexual interest; sigmoid displays; time to the eyed ova stage. |
|  |  | Fish and aquatic invertebrates | Bao et al. 2025 | Total number of eggs; offspring mortality; offspring length; offspring weight; hatching rate; offspring indicators (weight, mortality, malformation rate, and length); gonadosomatic index; time to emergence; offspring hatching rate; testosterone content; estradiol content; reproduction-related gene expression (vg1 and vtg2). |
|  |  | Fish and aquatic invertebrates | Berlino et al. 2023 | Oocyte production; sperm motility; brood size; embryo production; egg production; fertilisation rate. |
|  |  | Fish and aquatic invertebrates | Moyo et al. 2022 | Fecundity; gonad size; abnormal offspring (%); pregnancy probability. |
|  |  | Aquatic invertebrates | Funke et al. 2024 | Number of offspring. |
|  |  | Aquatic invertebrates | Mason et al. 2022 | Reproductive success; live young (%); sperm velocity; oocyte number; fecundity. |
|  |  | Terrestrial invertebrates | Ji et al. 2021 | Brood size; number of offspring. |
| **Survival** | Mortality | Fish | Sacco et al. 2026 | Number of deaths in each experimental group. |
|  | Survival | Fish | Hossain & Olden 2022 | Survival percentage. |
|  |  | Fish | Salerno et al. 2021 | Survival rate; survival percentage. |
|  |  | Fish | Wang et al. 2024 | Mortality; survival rate. |
|  |  | Fish | Yuan et al. 2025 | Maximum lifespan; mean lifespan; mortality; survival rate. |
|  |  | Fish and aquatic invertebrates | Berlino et al. 2023 | Survival rate; survival percentage. |
|  |  | Fish and aquatic invertebrates | Moyo et al. 2022 | Longevity; survival number; survival rate. |
|  |  | Aquatic invertebrates | Mason et al. 2022 | Mortality rate; survival rate; number of live individuals (%). |
|  |  | Terrestrial invertebrates | Abbas et al. 2025 | Survival. |
|  |  | Terrestrial invertebrates | Ji et al. 2021 | Number alive at the end of the experiment. |
|  |  | Terrestrial invertebrates | Li et al. 2024 | Survival rate. |

**Figure S2**

A summary of natural log response ratio (lnRR) effect-size estimates across 3 of the 14 included meta-analyses, comprising vertebrates (k = 356) and terrestrial invertebrates (k = 534). Natural log response ratio (lnRR) effect-size estimates ±95% confidence intervals (CI) for eight biological response trait categories under microplastic exposure: A) behaviour & locomotion, B) feeding & ingestion, C) growth & development, D) immunity, E) morphology, F) physiology & health, G) reproduction, and H) survival. Circles represent lnRR effect-size estimates and error bars represent 95% confidence intervals. The dashed vertical line indicates no effect (lnRR = 0). The y-axis shows the meta-analysis reference, the specific response variable extracted from each meta-analysis, and the number of effect-size estimates (k). Taxonomic groups are indicated by the colour of the y-axis labels: fish (blue) and terrestrial invertebrates (orange). Point colours indicate the biological interpretation of the response relative to the control: negative (red) and neutral (grey). Positive effect sizes for some endpoints (e.g., intestinal health) therefore appear on the right-hand side but are coloured red because they represent increased adverse effects.

**Figure S3**

A summary of odds ratio (OR) effect-size estimates across 2 of the 14 included meta-analyses, comprising vertebrates (k = NA) and terrestrial invertebrates (k = 26). Odds ratio (OR) effect-size estimates ±95% confidence intervals (CI) for five biological response trait categories under plastic exposure: A) behaviour & locomotion, B) feeding & ingestion, C) growth & development, D) reproduction, and E) survival. Circles represent odds ratio (OR) effect-size estimates, and error bars represent 95% confidence intervals (CI). The x-axis is displayed on a logarithmic scale. The dashed vertical line indicates no association (OR = 1), and confidence intervals overlapping one indicate non-significant associations. Odds ratios >1 indicate increased odds of the measured outcome, whereas odds ratios <1 indicate decreased odds of the measured outcome. The y-axis shows the meta-analysis reference, the specific response variable extracted from each meta-analysis, and the number of effect-size estimates (k). Taxonomic groups are indicated by the colour of the y-axis labels: fish (blue) and terrestrial invertebrates (orange). Point colours indicate the biological interpretation of the response relative to the control: negative (red) and neutral (grey). Positive effect sizes for some endpoints (e.g., mobility) therefore appear on the right-hand side but are coloured red because they represent increased adverse effects.
