## Supplementary figures and images for "Beyond harm: synthesising meta-analytical and mechanistic evidence on animal responses to microplastics"

### Figure S2

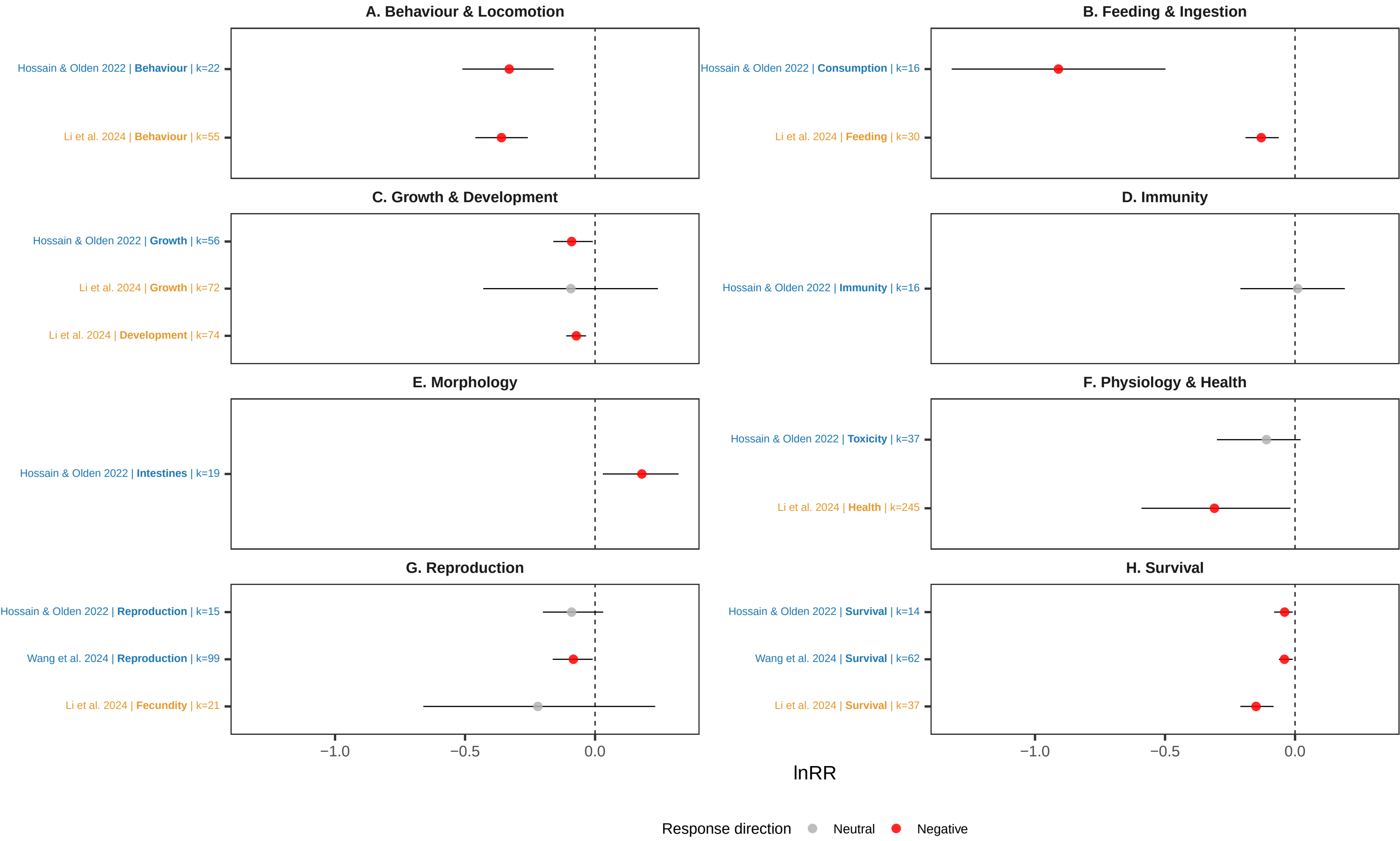

### Figure S3

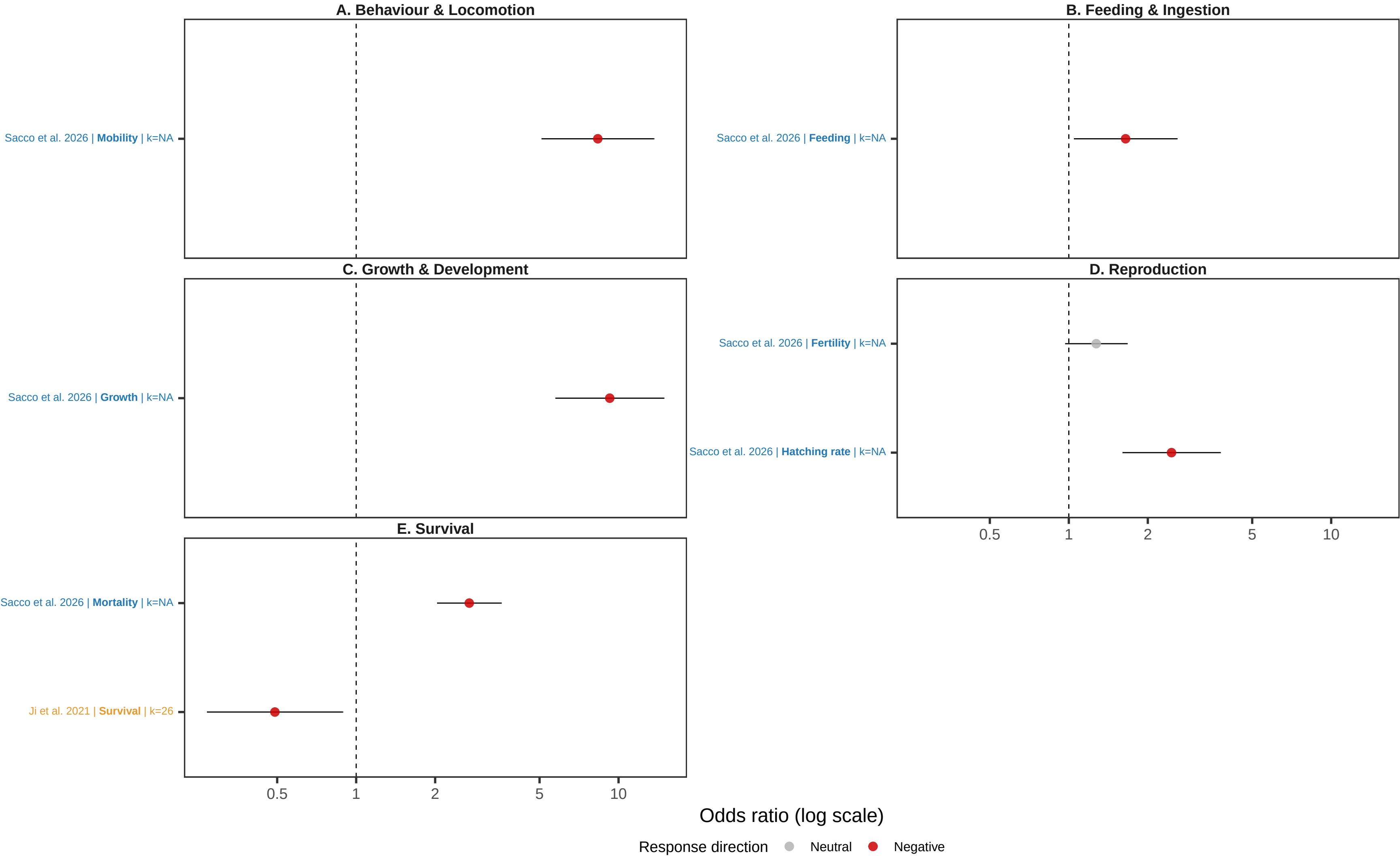
